# Functional genetic characterization of stress tolerance and biofilm formation in *Nakaseomyces glabrata* via a novel CRISPR activation system

**DOI:** 10.1101/2023.12.07.569977

**Authors:** Laetitia Maroc, Hajer Shaker, Rebecca S. Shapiro

**Affiliations:** Department of Molecular and Cellular Biology, University of Guelph, Guelph, Canada, N1G 2W1

## Abstract

The overexpression of genes frequently arises in *Nakaseomyces* (formerly *Candida*) *glabrata* via gain-of-function mutations, gene duplication or aneuploidies, with important consequences on pathogenesis traits and antifungal drug resistance. This highlights the need to develop specific genetic tools to mimic and study genetic amplification in this important fungal pathogen. Here, we report the development, validation, and applications of the first CRISPR activation (CRISPRa) system in *N. glabrata* for targeted genetic overexpression. Using this system, we demonstrate the ability of CRISPRa to drive high levels of gene expression in *N. glabrata*, and further assess optimal guide RNA targeting for robust overexpression. We demonstrate the applications of CRISPRa to overexpress genes involved in fungal pathogenesis and drug resistance, and detect corresponding phenotypic alterations in these key traits, including the characterization of novel phenotypes. Finally, we capture strain variation using our CRISPRa system in two commonly used *N. glabrata* genetic backgrounds. Together, this tool will expand our capacity for functional genetic overexpression in this pathogen, with numerous possibilities for future applications.

## Introduction

Fungal infections are a significant and growing threat to human health, posing a unique clinical challenge as a result of severely limited diversity of antifungal drugs, and therefore a scarcity of treatment options once drug-resistant infections are diagnosed. Recently, the WHO developed the first fungal priority pathogens list (1), highlighting the critical impact of fungal disease on human health, and emphasizing antifungal resistance as a “top priority” (1). Amongst the WHO high-priority pathogens is *Nakaseomyces* (formerly *Candida*) *glabrata*: an opportunistic yeast pathogen of emerging concern. *N. glabrata* is the second most common cause of candidiasis infections, accounting for ∼15-25% of invasive infections (2–4). While *Candida albicans* remains the most common cause of candidiasis, *N. glabrata*, along with other ‘non-*albicans Candida*’ species have been increasing in overall prevalence due in part to reduced susceptibility to antifungal drugs. Indeed, ∼20-30% of *N. glabrata* isolates are resistant to azole antifungals, especially fluconazole (5–10), and resistance to the echinocandin antifungals has increased significantly (from 5 to 12% of isolates) in the past decades (4, 9) with ∼15% of fluconazole-resistant isolates exhibiting multi-drug resistance (4). Thus *N. glabrata* is a critical pathogen with limited therapeutic options requiring focused research efforts to characterize its pathogenicity and mechanisms of antifungal resistance.

While numerous molecular mechanisms are involved in *N. glabrata*’s ability to resist treatment with antifungal agents, a common strategy involves the overexpression of factors involved in enabling growth in the presence of these drugs. Overexpression of drug efflux pumps belonging to both the ATP Binding Cassette (ABC) superfamily (including *CDR1* and *CDR2*) and the Major Facilitator Superfamily (MFS) classes is a common strategy to enable *N. glabrata* to remove antifungals from the cell (4, 7, 11). Antifungal-resistant *N. glabrata* strains are commonly identified with gain-of-function mutations in the transactional regulator Pdr1, which promotes overexpression of ABC transporters (7, 12–17). Further, *ERG11*, which encodes the target of the azole antifungals, is found to be upregulated upon azole treatment in *N. glabrata* and other *Candida* species (18), and the *ERG11* gene has been found in increased copy number in resistant isolates, due to duplication of the *ERG11*-containing chromosome (19–21). In addition to an important role in antifungal drug resistance, gene overexpression or amplification in *N. glabrata* is also associated with changes in virulence and interactions with the environment and host species (22–24), highlighting the critical role of genetic overexpression in *N. glabrata* adaptation.

This critical role for genetic overexpression in numerous facets of *N. glabrata* biology demands the development of tools to functionally characterize the overexpression of genes in this important pathogen. Currently, there are several genetic strategies for overexpression in *N. glabrata*, including promoter replacement strategies to drive high levels of expression (25, 26) and engineered expression plasmids (27). These techniques have provided important advances for the study of *N. glabrata* and have successfully probed the function of transporter genes involved in antifungal drug resistance. However, there are also limitations to the existing systems, which may rely on poorly efficient homologous recombination for integration, or time-consuming cloning or expensive synthesis of large plasmid constructs. Thus new techniques, including ones developed in other model systems (28) or well-studied *Candida* species (29), may be applied in *N. glabrata* to bolster the currently available genetic tools and expand our capacity for functional gene overexpression in this pathogen.

One advance that has significantly facilitated our capacity for genetic manipulation across a breadth of fungal species (30–34), is CRISPR-Cas-based techniques. CRISPR activation (CRISPRa) is one such strategy, which relies on an endonuclease dead Cas protein (*e.g.* dCas9) fused to transcriptional activators. This dCas9-activator complex can be targeted to the promoter region of a target gene via CRISPR single guide RNAs (sgRNAs) in order to achieve efficient gene activation/overexpression from a gene of interest. CRISPRa has been applied widely in mammalian cell lines, *Saccharomyces cerevisiae*, and other model systems, and has exploited a diversity of transcriptional activators to achieve robust gene overexpression (35–38). More recently, CRISPRa techniques have also been applied in fungal pathogens and other fungal species. In *C. albicans*, CRISPRa techniques have been successfully applied to drive high levels of expression of genes involved in biofilm formation, antifungal drug resistance, and stress tolerance, and phenotypically validate their role in these processes (39–41).

CRISPRa techniques have also been applied in diverse filamentous fungi for the activation of biosynthetic gene clusters and enhanced production of key bioactive metabolites (42–45). While CRISPR techniques have been adapted for diverse applications in *N. glabrata* (30, 46–48), including a recent CRISPR interference (CRISPRi) technique using similar principles to CRISPRa for gene repression (49), to date no CRISPRa systems have been developed for *N. glabrata*.

Here, we demonstrate the development and first applications of a CRISPRa genetic overexpression system for the fungal pathogen *N. glabrata*. This episomal plasmid-based CRISPRa system exploits a nuclease-dead dCas9 fused to a tripartite activator complex VP64-p65-Rta (VPR), and a Gibson assembly cloning method for highly-efficient single guide RNA integration. We show that this system can drive high levels of genetic overexpression and recapitulate associated antifungal drug resistance and biofilm growth phenotypes. We describe optimal guide RNA targeting rules for robust overexpression in *N. glabrata* and characterize novel phenotypes involved in stress and drug resistance associated with overexpression.

Finally, we capture strain variation using our CRISPRa system in two commonly used *N. glabrata* genetic backgrounds. Together, this tool will expand our capacity for functional genetic overexpression in this pathogen with numerous possibilities of applications.

## Results

### Development of a single CRISPRa plasmid in *N. glabrata*

In order to develop a CRISPRa system for genetic overexpression in *N. glabrata*, we designed and constructed a single-plasmid CRISPRa plasmid optimized for targeted gene activation in this fungal organism (Figure 1a). The system relies on a unique self-replicating and curable plasmid which allows the reversible overexpression of any gene without the need for stable genome modification. In this system, the genome-targeting portion of the guide RNA, the CRISPR RNA (crRNA) sequences, are cloned under the control of the RNA polymerase III *SNR52p* promoter at the NotI-restriction site, which has been previously validated as a facile system for highly efficient Gibson assembly-based cloning of sgRNA targeting sequences (50). In the same plasmid, we cloned the *dCAS9* gene under the control of the strong and constitutive *N. glabrata PDC1* promoter (51) and fused *dCAS9* to the tri-domains VPR activator complex developed in *S. cerevisiae* (52). The VPR activator complex is composed of two SV40-NLS sequences, a VP64 domain (composed of four repeats of the minimal activation domain of herpes simplex virus VP16), a SV40-NLS sequence, a linker, a p65 domain (transcriptional activation domain of human RelA), and a Rta AD domain (transcriptional activation domain from the human herpesvirus 4 (Epstein-Barr virus) replication and transcription activator Rta/BRLF1). The VPR activator complex fused to *dCAS9* has previously been demonstrated to robustly activate gene expression in both *S. cerevisiae* and *C. albicans* (*39, 52*). This newly-developed *N. glabrata* CRISPRa plasmid has been deposited to Addgene (Supplementary Table 1, Addgene plasmid catalog #213041).

**Figure 1.**
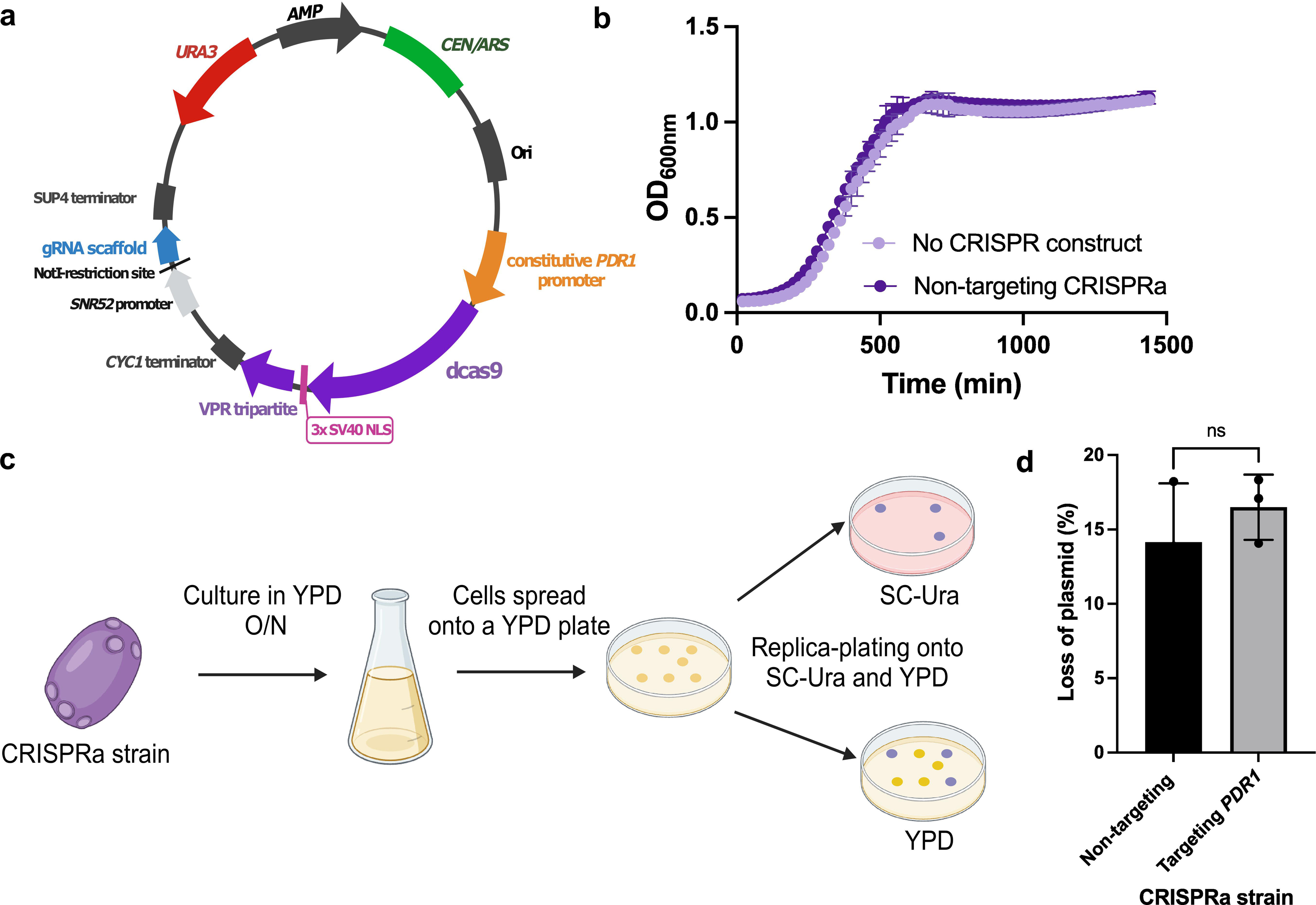
The *Nakaseomyces glabrata* CRISPRa system. (a) Plasmid map of *N. glabrata* CRISPRa plasmid (pCGLM2 aka pRS712). pCGLM2 is a centromeric Ura+ selectable plasmid. The catalytically dead Cas9 (dCas9) is fused to the VPR activator complex, which is composed of the VP64, p65 and Rta AD activator domains. sgRNAs are cloned at the NotI-restriction site using Gibson assembly. Panel created with BioRender.com, not to scale. (b) Growth of the CRISPRa backbone plasmid (no dCas9-VPR or sgRNA construct) compared with growth of a strain containing a non-targeting CRISPRa plasmid. Cells were grown in SC-Ura at 37°C. Growth was monitored by Optical Density (OD) reading at 600 nm every 20 min for 24 hrs. (c) Curing of the CRISPRa plasmid from *N. glabrata*. Cells that have lost the CRISPRa plasmid are selected after two rounds of culture in YPD at 37°C. ns: not significant (*p* value>0.05), Wilcoxon-Mann-Whitney Utest. All experiments were carried out in three independent replicates.

To confirm that the dCas9-VPR construct did not impact *N. glabrata* growth, we compared the growth of a strain transformed with the backbone plasmid, pCU-PDC1, from which the CRISPRa plasmid was constructed, with a strain transformed with the CRISPRa plasmid encoding a non-targeting sgRNA. We found no significant impact of the dCas9-VPR construct on *N. glabrata*’s growth (Figure 1b), suggesting the CRISPRa plasmid does not significantly impact fitness under the conditions tested.

Yeast episomal plasmids have the advantage of being curable from cells by removing the selection pressure, making any system based on these vectors fully reversible. Since our system relies on the *URA3* selection marker, cells that have lost the plasmid should be easily selectable after culturing in a non-selective medium, (i.e., in YPD). To demonstrate that our CRISPRa plasmid can be cured from *N. glabrata* cells, we cultured a strain transformed with a non-targeting CRISPRa and a strain transformed with the CRISPRa plasmid overexpressing *PDR1*. We found that our CRISPRa plasmid can be easily cured from cells after two rounds of culture in YPD with a successful rate of ∼14-16%, and that whether or not the system is actively targeting a gene does not impact this curability (Figure 1c).

### Implementation of CRISPRa as a tool for gene activation in N. glabrata

#### Overexpressing a gene implicated in fluconazole resistance

To validate that this CRISPRa system could effectively overexpress genes in *N. glabrata*, we first targeted a gene with a well-characterized antifungal drug resistance phenotype associated with high levels of expression. We focused on *PDR1*, encoding a transcriptional regulator of a pleiotropic drug resistance network in *N. glabrata,* whose overexpression is known to play a role in resistance to azole antifungals in clinical isolates (16, 53–55). Since previous research reported different transcriptional start sites (TSS) for *N. glabrata*’s *PDR1* gene (56, 57), we decided to design and clone 10 unique sgRNAs for our CRISPRa plasmid, spanning the promoter region of *PDR1* from -736bp to +36bp (relative to the start codon) on both sense and antisense DNA strands. We then determined the susceptibility of these CRISPRa strains in fluconazole using a Minimum inhibitory concentration (MIC) assay (Figure 2a). We were able to see a decrease in fluconazole susceptibility (MIC50 shifted from 80 µg/mL to 160 µg/mL) with sgRNAs located at -553 bp and -550 bp, on the sense and antisense strands respectively.

**Figure 2.**
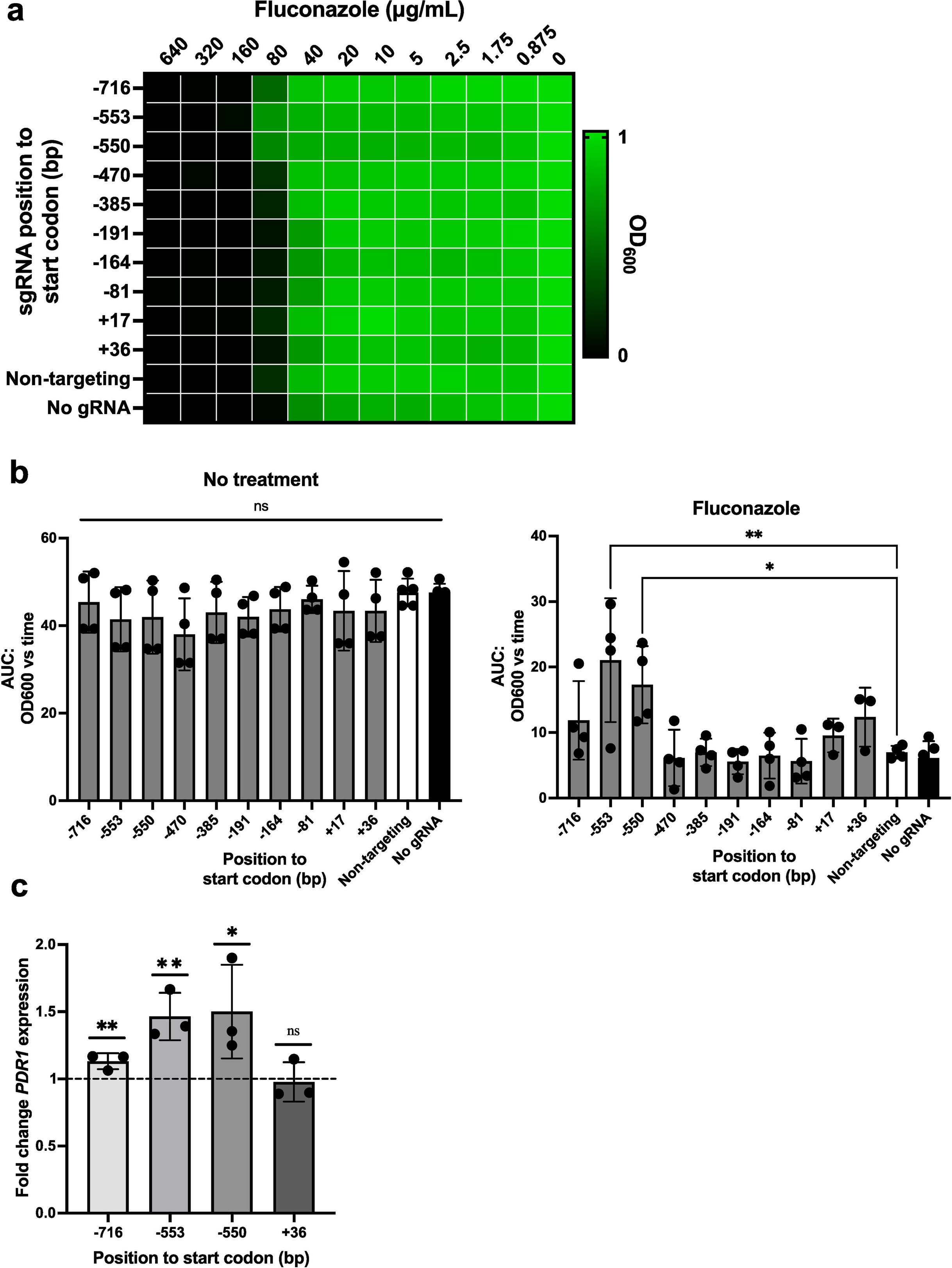
Validation of the *N. glabrata* CRISPRa system for overexpressing a gene involved in fluconazole resistance. (a) The fitness of CRISPRa strains overexpressing *PDR1* was measured using a MIC assay. Results are presented as a heatmap in which the growth in drugs, relative to the growth in no drug, is depicted, with the value 1.0 representing the highest fitness. (b) The fitness of CRISPRa strains overexpressing *PDR1* was measured using growth curves assays in the absence of drug (no treatment) and in 80 µg/mL of fluconazole. The relative area under the time-versus-optical dentistry (OD) curve (AUC) was then calculated. Dots represent the mean AUC and errors bars the standard deviation. Differences between groups were tested for significance using a Kurskal-Wallis test for the no treatment condition and a one-way ANOVA for the fluconazole condition. ns: not significant (*p* value>0.05); “*”: *p* value<0.05; “**”: *p* value<0.01. All experiments were carried out at least three times independently. (c) Fold change expression of *PDR1* in CRISPRa strains overexpressing *PDR1*. Fold change in the expression of *PDR1* relative to the housekeeping gene 18S rRNA in both experimental and non-targeting strains was measured by RT-qPCRs. Relative expression was then calculated using the non-targeting CRISPRa strain and the mean fold differences were plotted. Error bars depict the standard deviation. The dashed line represents the normalized baseline expression of the control strain. The fold change differences were tested for significance using a Student’s T-test. ns: not significant (*p* value>0.05); “*”: *p* value<0.05; “**”: *p* value<0.01. All experiments were carried out in at least three independent replicates.

To further validate the decreased susceptibility in our CRISPRa strains, we performed growth curve assays in media containing 80 µg/mL of fluconazole and calculated the relative area under the curve (AUC). As a control, strains were also cultured in media without drugs (Figure 2b). No fitness differences between strains were observed when strains were cultured without drugs, however, in fluconazole, the two CRISPRa strains encoding sgRNAs located at - 553 bp and -550 bp were significantly increased in their ability to grow compared to the non-targeting CRISPRa strain.

To confirm phenotypic changes in drug susceptibility were due to transcriptional overexpression of *PDR1*, we quantified the expression of *PDR1* in four different CRISPRa strains via RT-qPCR compared with the control strain encoding a non-targeting gRNA (Figure 2c). We found that our CRISPRa system enhances *PDR1* transcription by ∼1.5 fold and that this was achieved by using sgRNAs targeting both the sense or antisense DNA strands. Together, this indicates that our CRISPRa system can drive overexpression of genes in *N. glabrata*, and recapitulate phenotypes associated with genetic overexpression.

#### Overexpressing a gene implicated in biofilm formation

To demonstrate that our CRISPRa system can be used to assess genes involved in fungal pathogenesis traits, we decided to overexpress *EFG1*, encoding a transcription factor implicated in the control of biofilm formation (58). This gene has been characterized in *N. glabrata* and is known to lead to enhanced biofilm growth when overexpressed (58). Since the TSS of *EFG1* is well-defined, we were able to design 10 sgRNAs to span the promoter of *EFG1*. We generated 10 *N. glabrata* strains each containing unique sgRNAs located –462 to +112bp relative to the TSS, and measured the resultant biofilm growth of these strains using XTT-reduction biofilm assays. Three of these CRISPRa strains (encoding gRNAs located at -190 bp, -105 bp and -36 bp) significantly have a better ability to form biofilms as compared to the non-targeting CRISPRa strain (Figure 3a).

**Figure 3.**
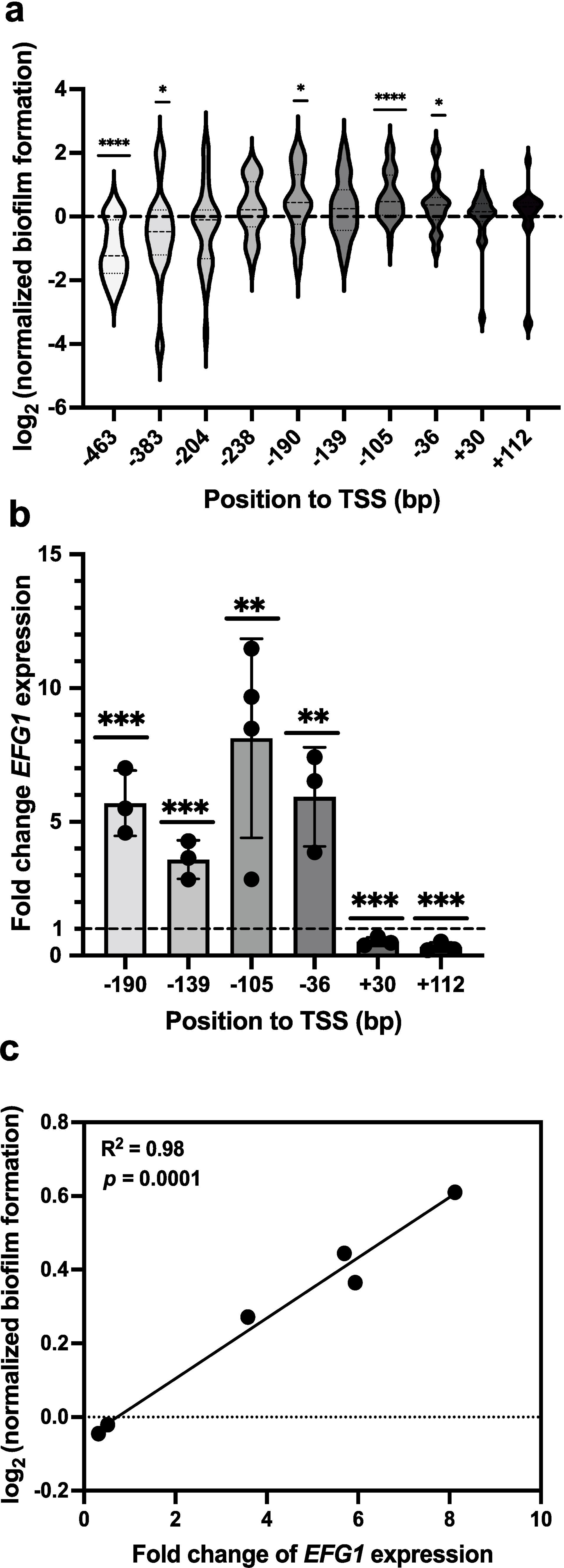
Validation of the *N. glabrata* CRISPRa system for overexpressing a gene involved in biofilm formation. (a) The biofilm-forming capacity of CRISPRa strains overexpressing *EFG1* was measured using an XTT-reduction biofilm assay. The relative biofilm formation of these strains was calculated using the non-targeting strain as a control and the relative log_2_ values were calculated. Long dashes on violin plots represent the median and short dashes are the first and third quartiles. Data were combined from three independent experiments where n = 20 for +30 and +112 sgRNAs encoding strains and n=32 for the remaining strains. Differences between groups were tested for significance using a one-sample Wilcoxon test. ns: not significant (p-value>0.05); “*”: *p* value<0.05; “***”: *p* value<0.001: “****”: *p* value<0.0001. (b) Fold change expression of *EFG1* in CRISPRa strains overexpressing *EFG1*. Fold change in the expression of *EFG1* relative to the housekeeping gene 18S rRNA in both experimental and non-targeting strains was measured by RT-qPCRs. Relative expression was then calculated using the non-targeting CRISPRa strain and the mean fold differences were plotted. Error bars depict the standard deviation. The fold change differences were tested for significance using a Student’s T-test. “**”: *p* value<0.01; “***”: *p* value<0.001. All experiments were carried out in at least three independent replicates. (c) The square Pearson’s correlation coefficient between the biofilm capacity and the fold change in *EFG1* expression in the CRISPRa strains overexpressing *EFG1*.

We next profiled the expression of *EFG1* in our CRISPRa strains and found that *EFG1* was overexpressed from ∼3.6 fold to 8.1 fold using our dCas9-VPR construct, with the highest level of expression obtained when targeting the dCas9-VPR construct -105 bp upstream the TSS of *EFG1* (Figure 3b). Compellingly, we found a significant correlation between the level of *EFG1* overexpression and the biofilm-forming capacity of the CRISPRa strains (Figure 3c). We further found that targeting the CDS leads to repression of *EFG1*, which may be a result of dCas9-VPR sterically interfering with the RNA polymerase II (Figures 3b and 3c).

### Characterization of two new genes whose overexpression leads to decreased caspofungin susceptibility and better stress tolerance

Finally, we wanted to demonstrate that our CRISPRa system could be exploited to characterize new genes whose overexpression has not previously been profiled. For this, we targeted *N. glabrata STE11* and *SLT2* genes for overexpression. *STE11* encodes a mitogen-activated protein kinase and has been shown to be involved in control of stress response and virulence in *N. glabrata* (59). Notably, Ste11 has been shown to mediate cross-tolerance to different environmental stresses such as low pH or oxidative stress (60). In the same study, authors stipulated that the identified mutations in *STE11* were likely to enhance the activity of Ste11 but the role of *STE11*’s expression levels in stress tolerance has never been studied. *SLT2* is also predicted to encode a mitogen-activated protein kinase, but with a role in cell wall integrity (61). It has been shown that in *N. glabrata,* this gene becomes overexpressed following a caspofungin treatment (62), but no study has shown that the overexpression of *SLT2* is correlated to better tolerance to caspofungin.

We generated CRISPRa strains overexpressing *STE11* and *SLT2* ∼1.2-1.8 and ∼1.8-2.8 fold, respectively. We assessed thermotolerance and tolerance to H_2_O_2_-induced oxidative stress for the *STE11*-overexpressing CRISPRa strains and caspofungin susceptibility for the *SLT2*-overexpressing ones. We were able to demonstrate that overexpression of *STE11* and *SLT2* leads to a better tolerance to heat and oxidative stress, and to a better fitness in caspofungin, respectively. This demonstrates our ability to exploit this CRISPRa system to investigate novel phenotypes associated with gene overexpression in *N. glabrata*.

### The CRISPRa platform is functional in multiple *N. glabrata* strain backgrounds

General physiology differs across different strain backgrounds and, notably, it has been reported how stress sensitivity and growth physiology traits can vary significantly across yeast strains (63, 64). In *S. cerevisiae*, the use of CRISPR technologies across different strain backgrounds can significantly impact phenotypes, based on the genetic background (65). A recent study has demonstrated that phenotypic differences, notably in metabolic profiles and cell wall carbohydrates complexity and abundance, exist between the two commonly used *N. glabrata* isolates, CBS138 and BG2, and that it can influence its pathogenicity (66). All experiments described above in this study were carried out in strain HM100, derived from the reference strain CBS138. To determine possible strain variation responses to our CRISRa system, we decided to test our system in another *N. glabrata* strain background, BG87 derived from *N. glabrata* strain BG2. To do so, we overexpressed *PDR1* and *EFG1* in BG87, employing the same sgRNAs as those used in the HM100 strain.

We found that our CRISPRa system works efficiently in strain BG87, but that its portability is influenced by the strain. Indeed, we were able to overexpress *PDR1* and *EFG1* up to ∼1.8 and 5.7 fold, respectively (Figure 5a), however, of the two sgRNAs leading to *PDR1* overexpression in strain HM100, only one increased the transcriptional activity of *PDR1* and decreased FLZ susceptibility in strain BG87 (Figure 5b,c). Similar results were obtained for *EFG1*, where only one of three gRNAs that led to strong *EFG1* overexpression and a high capacity to form biofilm in strain HM100 increased strain BG87’s capacity to form biofilm (Figure 5d). These results may be due to TSS differences between the two strains.

**Figure 4.**
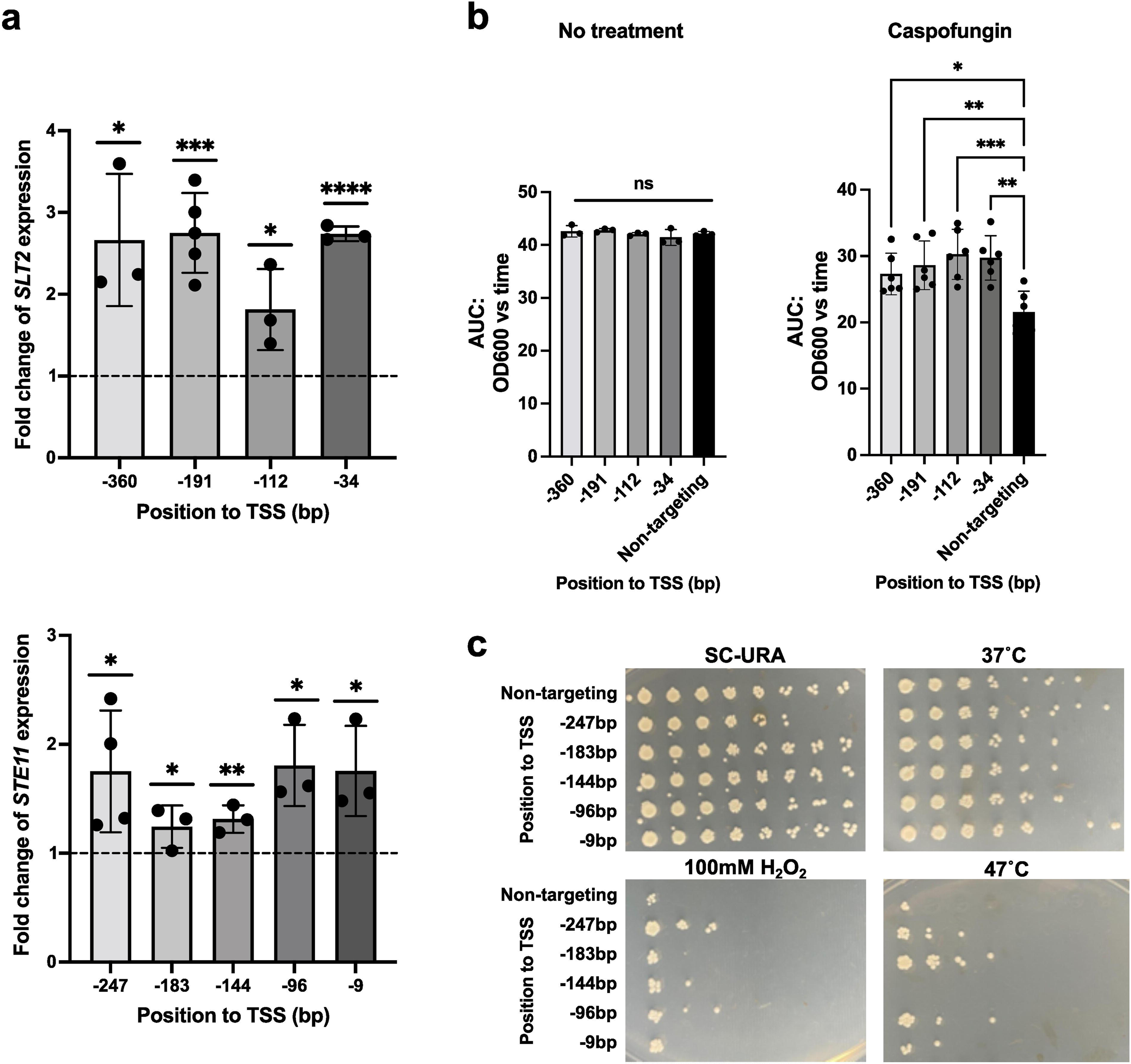
Characterization of *SLT2* and *STE11*’s overexpression in caspofungin susceptibility and stress tolerance. (a) Fold change expression of *SLT2* (upper panel) and *STE11* (lower panel) in CRISPRa strains overexpressing these genes. Fold change in the expression of *SLT2* and *STE11* relative to the housekeeping gene 18S rRNA in both experimental and non-targeting strains was measured by RT-qPCRs. Relative expression was then calculated using the non-targeting CRISPRa strain and the mean fold differences were plotted. Error bars depict the standard deviation. The fold change differences were tested for significance using a Student’s T-test. “*”: *p* value<0.05; “***”: *p* value<0.001; “****”: *p* value<0.0001. All experiments were carried out in at least three independent replicates. (b) The fitness of CRISPRa strains overexpressing *SLT2* was measured using growth curves assays in the absence of drug (no treatment) and in 32 ng/mL of caspofungin. The relative area under the time-versus-optical dentistry (OD) curve (AUC) was then calculated. Dots represent the mean AUC and errors bars the standard deviation. Differences between groups were tested for significance using a one-way ANOVA. ns: not significant (*p* value>0.05); “*”: *p* value<0.05; “**”: *p* value<0.01; “***”: *p* value<0.001. All experiments were carried out in at least three independent replicates. (c) Assessing stress tolerance in CRISPRa strains overexpressing *STE11* using serial dilutions spot assays. 10-fold serial dilutions of cells were spotted on SC-Ura after having undergone different treatments. Photographies were taken after cells spent 48 hrs at 37°C.

**Figure 5.**
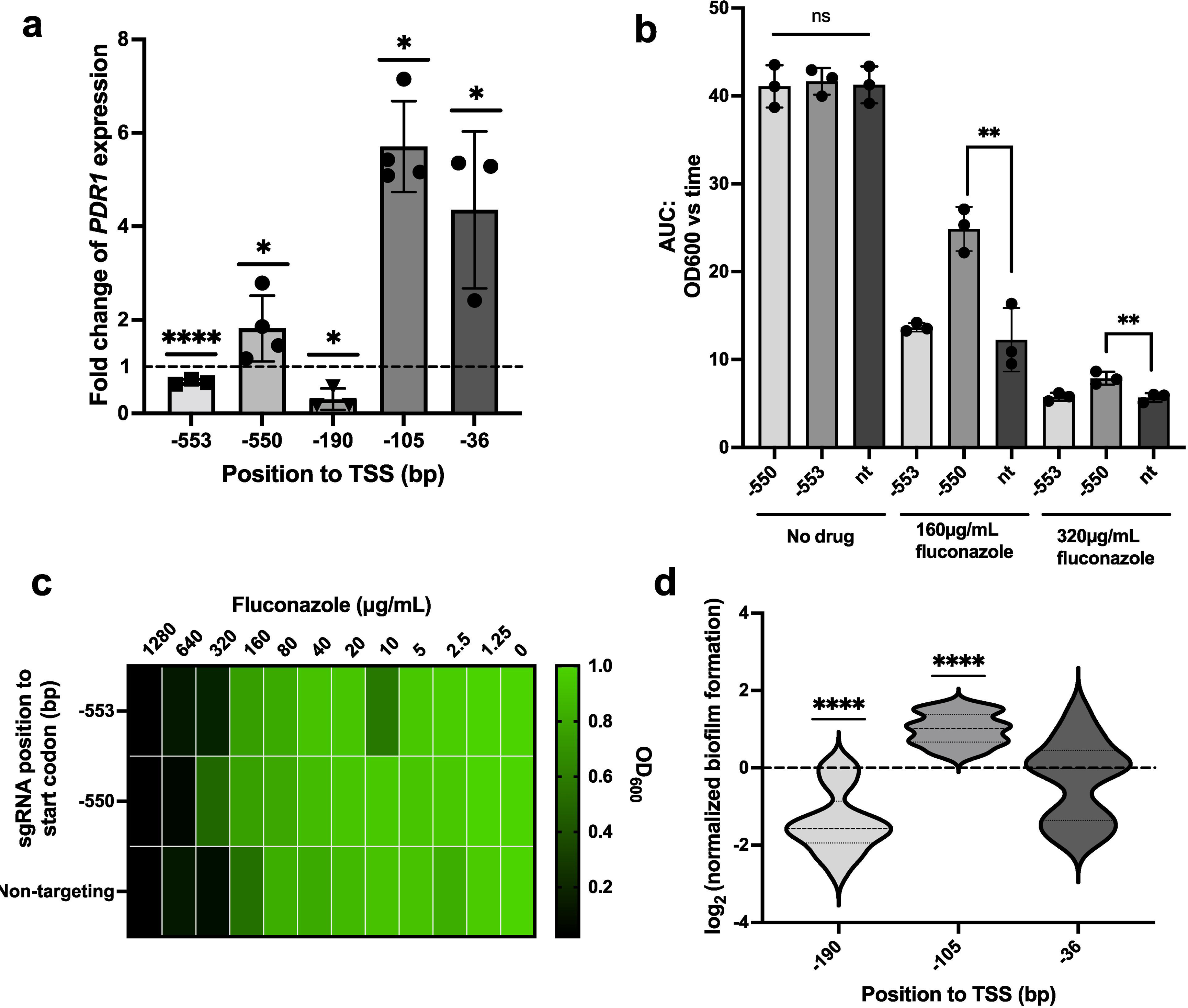
Validation of the *N. glabrata* CRISPRa system in strain BG87. (a) The fitness of CRISPRa strains overexpressing *PDR1* was measured using MIC assays. Results are presented as a heatmap in which the growth in drugs, relative to the growth in no drug, is depicted, with the value 1.0 representing the highest fitness. (b) The fitness of CRISPRa strains overexpressing *SLT2* was measured using growth curves assays in the absence of drug (no treatment) and in 160 or 320 µg/mL of fluconazole. The relative area under the time-versus-optical dentistry (OD) curve (AUC) was then calculated. Dots represent the mean AUC and errors bars the standard deviation. Differences between groups were tested for significance using a one-way ANOVA. ns: not significant (*p* value>0.05); “**”: *p* value<0.01. All experiments were carried out in at least three independent replicates. (c) The biofilm-forming capacity of CRISPRa strains overexpressing *EFG1* was measured using an XTT-reduction biofilm assay. The relative biofilm formation of CRISPRa strains overexpressing *EFG1* was calculated using the non-targeting strain as a control, and the relative log_2_ values were calculated. Long dashes on violin plots represent the median and short dashes are the first and third quartiles. Data were combined from three independent experiments where n = 21. Differences between groups were tested for significance using a one-sample Student’s T-test. “****”: *p* value<0.0001. (d) Fold change expression of *PDR1* and *EFG1* in CRISPRa strains overexpressing *PDR1* and *EFG1*, respectively. Fold change in the expression of *PDR1* and *EFG1* relative to the housekeeping gene 18S rRNA in both experimental and non-targeting strains was measured by RT-qPCRs. Relative expression was then calculated using the non-targeting CRISPRa strain and the mean fold differences were plotted. Error bars depict the standard deviation. The fold change differences were tested for significance using a Student’s T-test: “*”: *p* value<0.05; “****”: *p* value<0.0001. All experiments were carried out in at least three independent replicates.

## Discussion

Here, we report the development, validation and application of the first CRISPRa technology in *N. glabrata.* Where nearly all CRISPR systems developed in *N. glabrata* rely on dual plasmid systems (46, 49), ours relies on a single centromeric plasmid which limits the need for selectable markers, and is not reliant on inefficient homologous recombination in *N. glabrata,* as no stable genome integration is needed. Since it is a centromeric plasmid, selection pressure must be maintained during experiments in order to maintain plasmid-containing cells. Our CRISPRa system further employs *URA3* as a selectable marker, which allows a non-toxic selection pressure and avoids the risk of potential synergistic effects with other drugs used during experimentimentation. In our plasmid system, sgRNAs can easily be cloned at the NotI-restriction site using the highly efficient Gibson assembly method. Finally, since our system is constitutive, it does not require specific inducing conditions and is active as soon as it is transformed into cells which facilitates its use.

We have demonstrated the ability of our CRISPRa system to increase the transcriptional activity of *N. glabrata*’s genes ∼1.5 to 8 fold. We found that designing and testing only ∼2-3 sgRNAs in regions -360 to -9 bp upstream of the TSS of the target gene on either DNA strand is sufficient to achieve high levels of overexpression. The ability to target both the sense and antisense DNA strands by our dCas9-VPR construct increases the number available PAM sites, and thus increases the likelihood of identifying usable sgRNAs. Finally, we find that the level of overexpression achieved can be conveniently regulated based on the targeting location of the sgRNA within the promoter region of the target gene, which effectively makes the system titratable. This has been previously reported in many CRISPRa studies in other organisms (35, 39, 40, 67), and constitutes a powerful feature of this and other CRISPRa technologies.

Interestingly, for two genes, we found that a slight increase in transcriptional activity (∼1.46-1.50 fold for *PDR1* and 1.24-1.80 fold for *STE11*) was enough to lead to detectable phenotypes: decreased susceptibility to fluconazole for *PDR1* and better stress tolerance for *STE11*. This is in contrast to *EFG1*, where expression needed to be increased over 3.5 fold to promote enhanced biofilm growth. For *PDR1*, gain-of-function mutations/duplication of the *PDR1* gene or *PDR1*-containing chromosome is frequently identified as a mechanism of resistance in fluconazole-evolved laboratory and clinical isolates of *N. glabrata* (17, 53, 55, 68–71). While expression of *PDR1* in these strains has not typically been studied, our results suggest that even if mutations lead to a weak increased transcriptional activity (below 2-fold), it may lead to reduced susceptibility to fluconazole, which clinically represents a threat to treatment success. In addition, we found a strong correlation between the level of overexpression induced by our dCas9-VPR construct and the magnitude of the resulting phenotype. The ability to modulate the levels of expression using diverse sgRNAs, allows us to mimic dose-dependent phenotypes that are known to exist in *Candida* species, such as azole drug resistance associated with dose-dependent expression of *TAC1* and *ERG11* (72).

The validation of our system in two *N. glabrata* genetic backgrounds enables us to capture strain variation. Some sgRNAs lead to overexpression in one strain and to repression in another, possibly due to differing TSSs between the two strains. It is possible that the dCas9-VPR construct sterically prevents transcription factors from accessing their binding sites, therefore causing repression of the target gene. Using our CRISPRa system, we were also able to characterize new genes for which overexpression leads to a decrease in caspofungin susceptibility and better stress responsiveness, including thermotolerance. This underlies the importance of overexpression alteration in drug resistance and stress response in *N. glabrata*. While these genes have been previously studied, we demonstrate that novel forms of genetic alterations (i.e., deletions vs. overexpression, etc) can help characterize phenotypes in new ways, and therefore, we must use all available approaches (e.g, repression, activation, deletion) to fully capture the role and regulation of genes. This will ultimately help better predict and understand important facets of biology, including virulence and antifungal drug resistance, in an important fungal pathogen like *N. glabrata.* Altogether, this unique CRISPRa system will bolster the currently available genetic tools, and expand our capacity for functional gene overexpression in this pathogen. Applied at a genome-scale, this tool could lead to the identification of new genes implicated in drug resistance, virulence and stress response. This could lead to the identification of potential new drug targets to ultimately combat this opportunistic pathogen.

## Materials and methods

### Strains and growth conditions

Yeast strains, plasmids, and primers used in this study are listed in Table S1, Table S2 and Table S3, respectively. Yeast strains were grown in broth or on plates at 37°C in Synthetic Complete medium lacking uracil for plasmid selection (SC-Ura, 0.67% Yeast Nitrogen Base without amino acids with ammonium sulfate, 2% glucose, supplemented with adenine and all amino acids except uracil), YPD (2% peptone, 1% yeast extract and 2% glucose) or RPMI-1640 supplemented with 3.5 % morpholinepropanesulfonic acid (MOPS) and 2% glucose.

*Escherichia coli* DH5α cells were cultured at 30°C in broth or on LB agar plates containing 100 µg/mL ampicillin for plasmid selection.

### PCR, transformations and genomic DNA (gDNA) extraction

PCRs were performed using the Q5® High-Fidelity DNA Polymerase for cloning and crRNA N20 sequence amplification or FroggaBio Taq DNA Polymerase for transformants screening. gDNA was extracted using the Thermo Fisher Scientific Invitrogen™ PureLink™ Genomic DNA Mini Kit.

Yeast transformations were performed according to the “one-step” lithium acetate transformation protocol, as previously described (73). Bacterial transformations were performed using NEB® 5-alpha Competent *E. coli* cells, according to the manufacturer’s recommendations.

### Plasmid design and cloning

Molecular cloning was performed using the Gibson assembly method (74).The dCas9 backbone plasmid was synthesized by Twist Bioscience (www.twistbioscience.com) from plasmid pCU-MET3 (51). This dCas9 backbone plasmid contains the *URA3* selectable marker, an *N. glabrata* centromeric origin of replication (CEN/ARS sequence), sgRNA cloning site (*SNR52* promoter, NotI cloning locus, and sgRNA tail) and dCas9 (D10A mutation in the RuvC catalytic domain and N863A mutation in the HNH catalytic domain) cloned under the control of the *MET3* promoter (*MET3p*). The *MET3* promoter was replaced by the strong and constitutive *PDC1* promoter (*PDC1p*) to allow a strong expression of dCas9 (51). The endogenous *PDC1p* sequence was amplified using primers PDC1-F and PDC1-dCas9-R and gDNA of strain HM100 as a DNA template. Since primers PDC1-F and PDC1-dCas9-R contain 40 bp identical to regions surrounding the MET3p, it enables a replacement of *MET3p* by *PDC1p* into the dCas9 backbone plasmid by Gibson assembly. This gives rise to the PDC1-dCas9 plasmid. Into PDC1-dCas9, we fused a gene encoding dCas9 to a VPR three-part domain (2x SV40NLS - VP64 domain (composed of four repeats of the minimal activation domain of herpes simplex virus VP16) - 1xSV40 NLS-p65 domain (transcriptional activation domain of human RelA) - Rta AD (transcriptional activation domain from the human herpesvirus 4 (Epstein-Barr virus) replication and transcription activator Rta/BRLF1)) synthesized as a gene fragment gene from Twist Bioscience. The sequence of this gene fragment can be found in Supplementary Table 4.This gene fragment was cloned into the dCas9 plasmid backbone using Gibson assembly. The final construct (pCGLM2 aka pRS712) was fully sequenced by Plasmidsaurus (www.plasmidsaurus.com/) and is available via Addgene (www.addgene.org/; catalog number: #213041).

The control backbone plasmid, pCU-PDC1 was constructed from pCU-MET3 (51) by replacing the *MET3p* with *PDC1p*. The endogenous *PDC1p* sequence was amplified using primers PDC1-F and PDC1-R and gDNA from *N. glabrata* strain HM100 as a DNA template. Since primers PDC1-F and PDC1-R contain 40 bp identical to regions surrounding the *MET3p*, it enables a replacement of *MET3p* by PDC1p *into* the pCU-MET3 by Gibson assembly. This gives rise to the pCU-PDC1 plasmid.

### sgRNA design and cloning

The sgRNA CRISPR RNAs (crRNA; N20 nucleotide sequence complementary to the target genomic DNA) were designed based on efficiency and predicted specificity via the sgRNA design tool Eukaryotic Pathogen CRISPR gRNA Design Tool (EuPaGDT; http://grna.ctegd.uga.edu; (75)). Sequences and information on crRNA N20 sequences can be found in Supplementary Table 5. Each crRNA N20 sequence was ordered from Integrated DNA Technologies (www.idtdna.com) with 20 bp overlaps on either side of the region of target complementary and was amplified with extender oligos (primers gRNA-extender-F and gRNA-extender-R) to extend the overlaps from 20 to 40 bp, as previously described (50). They were then cloned at the NotI-restriction site into pCGLM2 using Gibson assembly.

sgRNAs were designed ∼ -716 to +36 bp upstream of the start codon for *PDR1* and ∼ - 463 to -9 bp from the TSS for *EFG1*, *STE11* and *SLT2*. Start codons and TSSs were obtained from The Candida Genome Database (CGD) (76).

### Growth curves

*N. glabrata* transformants carrying pCGLM2 were grown overnight in liquid SC-Ura medium at 37°C, 250 rpm, and used to inoculate fresh SC-Ura or SC-Ura containing either 80, 160 or 320 µg/mL of fluconazole (FLZ) or SC-Ura containing 32 ng/mL of caspofungin (CASP) at an OD_600_ of 0.05 or 0.005 in a 96-well flat-bottomed plate with a total volume of 200 µl per well topped with 25 µl of mineral oil to avoid evaporation. The starting OD_600_ of each strain was kept consistent within the same experiment. Growth was measured at 37°C via OD_600_ at 20 min intervals over the course of 24 h using an Infinite 200 PRO microplate reader (Tecan). Plates were shaken orbitally for 1,000 seconds at a 4.5-mm amplitude between growth measurements.

### Plasmid curing

CRISPRa strains were cultured overnight in YPD at 37°C , shaking at 250 rpm. The next day, cells were spread onto YPD plates to obtain single colonies and were grown for 48 hrs at 37°C. These plates were then replica-plated onto both SC-Ura and YPD plates and after 48 hrs at 37°C, cells that have lost the plasmid, growing onto YPD but not SC-Ura, were identified.

### Minimum inhibitory concentration (MIC) assays

MIC assays were performed in at least three replicates following the EUCAST E.DEF 7.3.2. protocol (77), with the following minor modifications. MICs were run in SC-Ura. Overnight cultures of *N. glabrata* grown in SC-Ura were diluted to an OD_530_ of 0.01 (∼2.10^5^ cells) in SC-Ura. Cells were mixed into the plates in an equal volume such that the starting OD_530_ values of each strain were 0.005 (∼0.5.10^5^ cells) in SC-Ura. Plates were incubated at 37°C without shaking and absorbance values at 530 nm were read after 24 h using an Infinite 200 PRO microplate reader (Tecan). The range of fluconazole concentrations tested was 1,280-1.25 µg/mL.

### Biofilm assays

Biofilm assays were performed as previously described (78), with minor modifications. *N. glabrata* cells from SC-Ura O/N cultures were diluted to an OD_600_ of 0.01 in RPMI in 200 µL in 96-well plates and cells were grown statically at 37°C for 48 hrs. After two washing steps in 1X PBS and once biofilms were dried, 90 µl of 1 mg/ml tetrazolium salt (XTT) (prepared in 1X PBS and centrifuged to remove sediment prior to use) and 10 µl of 0.32 mg/ml PMS (prepared in water) were added to each well. Plates were incubated statically at 30°C for 2 hrs to allow biofilms to reduce XTT, measured at 490 nm using an Infinite 200 PRO microplate reader (Tecan), and normalized to the growth of planktonic cells harvested previously. The “relative biofilm formation” was calculated as follows: (OD_490_/OD_600_ planktonic cells)_teststrain_/(OD_490_/OD_600_ planktonic cells)_control_, where the control represents the non-targeting CRISPRa strain.

### Stress assays

Stress assays were performed similarly to those previously described (60), with minor modifications. For the hydroxide peroxide (H_2_O_2_) stress assays, SC-Ura O/N cultures of the CRISPRa strains were diluted to an OD_600_ of 1 in a 96-deep well plate in 500 µL of SC-Ura or SC-Ura containing 100mM of H_2_O_2_. The plate was then incubated for 1 hr in a plate-shaking incubator at 37°C, shaking at 800 rpm. After incubation, cells were centrifuged in the 96-deep well plate at 300 g for 10 min to remove the supernatant. Cells were then resuspended in 100 µL of a sterile PBS solution, diluted in 10-fold serial dilutions in a PBS and each dilution was spotted using a 96-pin pinning tool onto SC-Ura. Plates were then photographed after 48 hrs at 37 °C.

For heat shocks, overnight cultures were diluted to an OD_600_ of 0.5 in SC-Ura in microcentrifuge tubes. Tubes on a tray were then incubated for 1 hr at 37°C or 47°C, statically. After the incubation, they were placed at 4°C for 5 min. Cells were then diluted and spotted on SC-Ura as described above. Plates were photographed after growth for 48 hrs at 37 °C.

### RNA extraction and real-time quantitative PCR (RT-qPCR)

To detect differences in gene expression, overnight cultures of *N. glabrata* grown in SC-Ura were diluted to an OD_600_ of 0.05 in fresh SC-Ura (for *SLT2* and *STE11*) or SC-Ura containing 80 or 160 µg/mL of FLZ (for *PDR1*) or RPMI (for *EFG1*) and grown to an OD_600_ of ∼0.6-0.8 at 37°C. Cells were pelleted and frozen at −80°C before RNA was extracted. RNAs were extracted using the RNeasy Mini kit from Qiagen, according to the manufacturer’s recommendations.

cDNA was synthesized from RNAs as follows: 1,000 µg of RNA were treated for DNA contamination using the Invitrogen™ TURBO DNA-free™ Kit from Thermo Fisher Scientific. The supernatant was then incubated for 5 min at 65°C with 4.5 µM of random hexamers (Thermo Fisher Scientific) and 9 µM of a dNTPs mix (Thermo Fisher Scientific). After the reaction, the mixture was put on ice for 1 min. From this cDNA was synthesized using the SuperScript™ IV Reverse Transcriptase from Thermo Fisher Scientific and was stored at - 20°C, as needed, before running RT-qPCRs.

RT-qPCRs were performed using a QuantStudio 3 Real-Time PCR Instrument (Thermo Fisher Scientific) with the SYBR Green-based method. Primers used for RT-qPCR are listed in Table S3. Expression profiling calculations were performed according to the comparative CT method (79). Briefly, expression values for the targeted gene of interest were compared with the housekeeping gene rRNA 18S to obtain a ΔCT value for each strain. The ΔCT values in the experimental CRISPRa strains were then compared with the non-targeting CRISPRa control strain to obtain a ΔΔCT value and a fold difference in the expression of the target gene.

### Analysis and statistics

All graphs and statistical analysis were performed using Graphpad Prism v9.4.1. Before using a parametric test, the normality of the datasets was checked. Normality was checked using the Shapiro-Wilk test.

## Acknowledgments

We thank P.r Cécile Fairhead for sharing strains HM100 and BG87. This work was supported by a Natural Sciences and Engineering Research Council of Canada (NSERC) Discovery Grant (RGPIN-2018-4914) and a Canadian Institutes for Health Research (CIHR) Project Grant (PJT 162195). RSS holds the Canada Research Chair (Tier II) in Microbial Functional Genomics. LM was supported by an EvoFunPath NSERC Collaborative Research and Training Experience (CREATE) fellowship.

## Supplementary material

**Supplementary Table 1.**
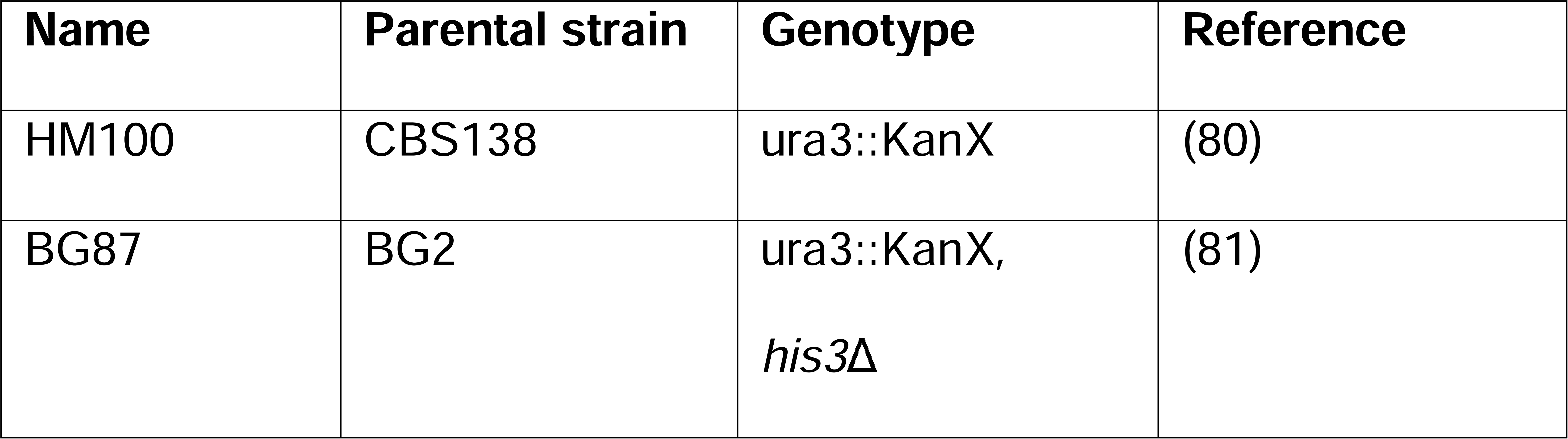
Strains used in this study.

**Supplementary Table 2.**
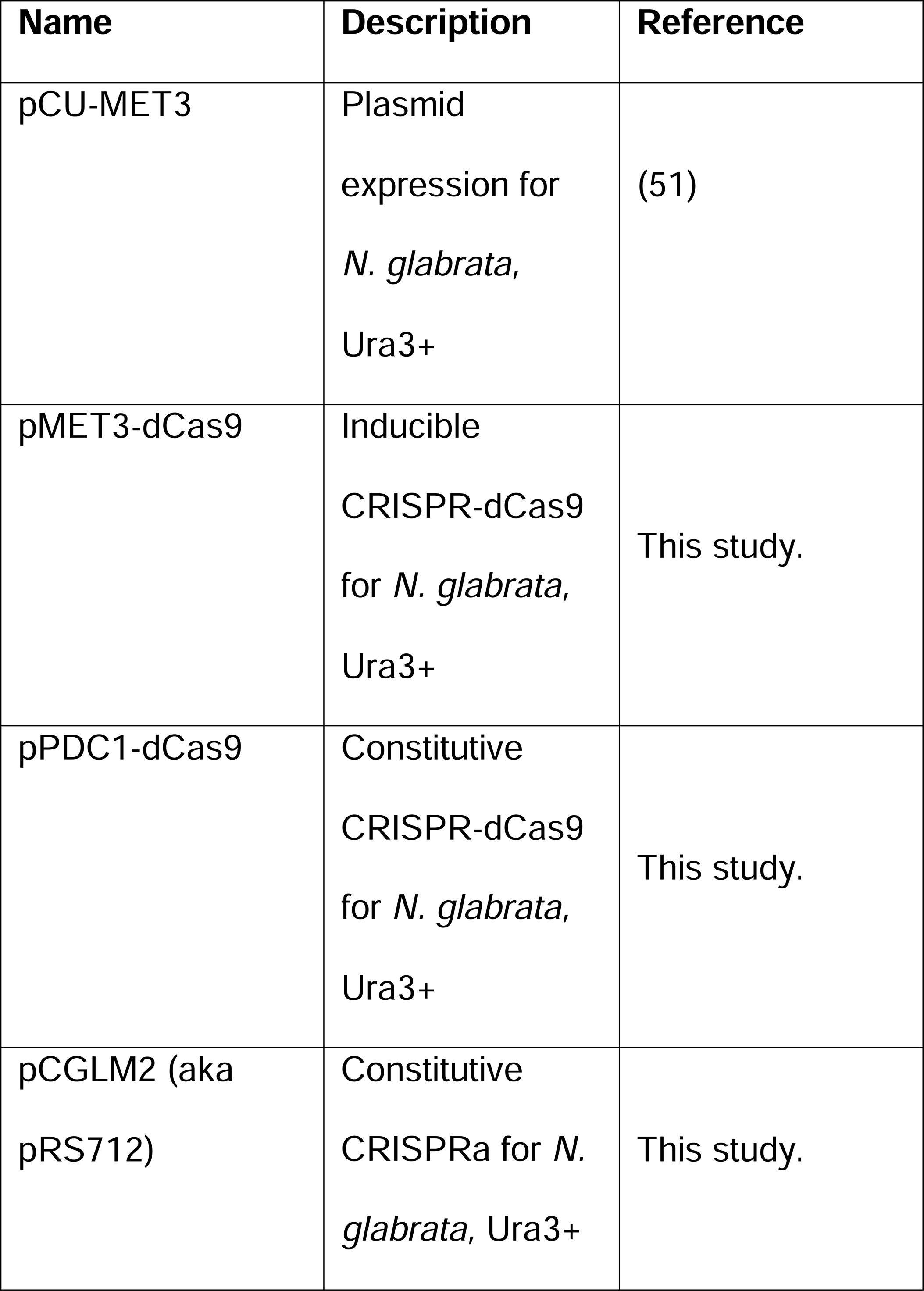
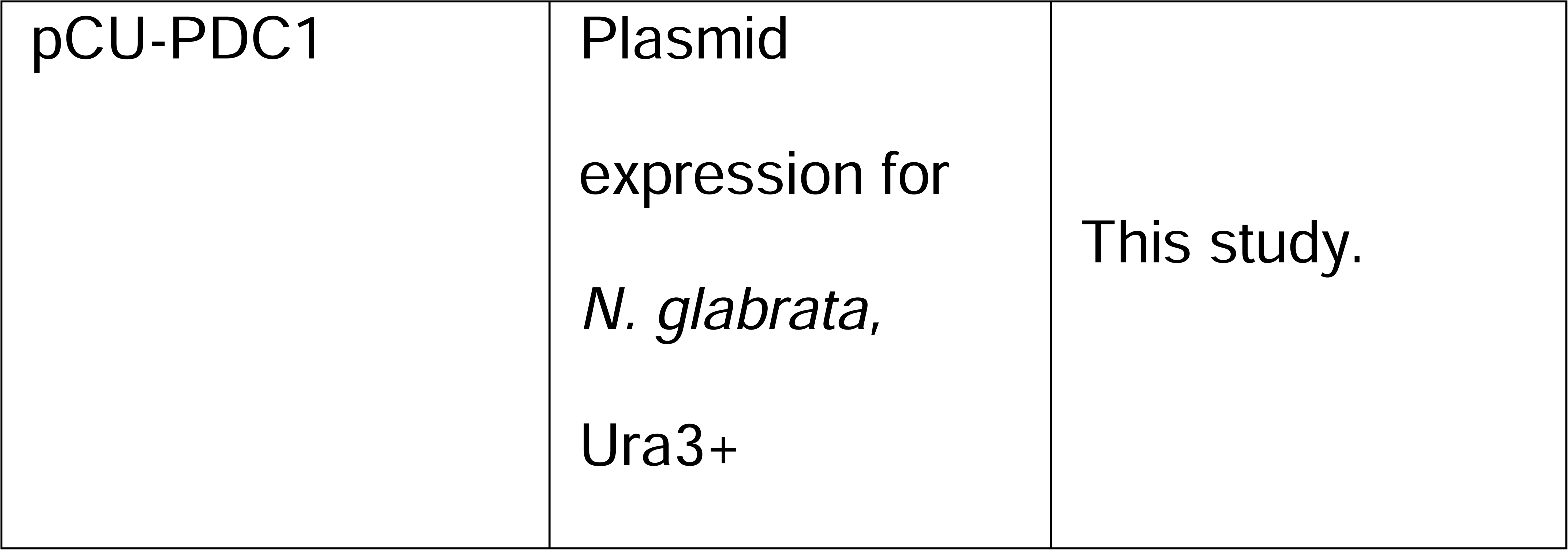
Plasmids used in this study.

**Supplementary Table 3.**
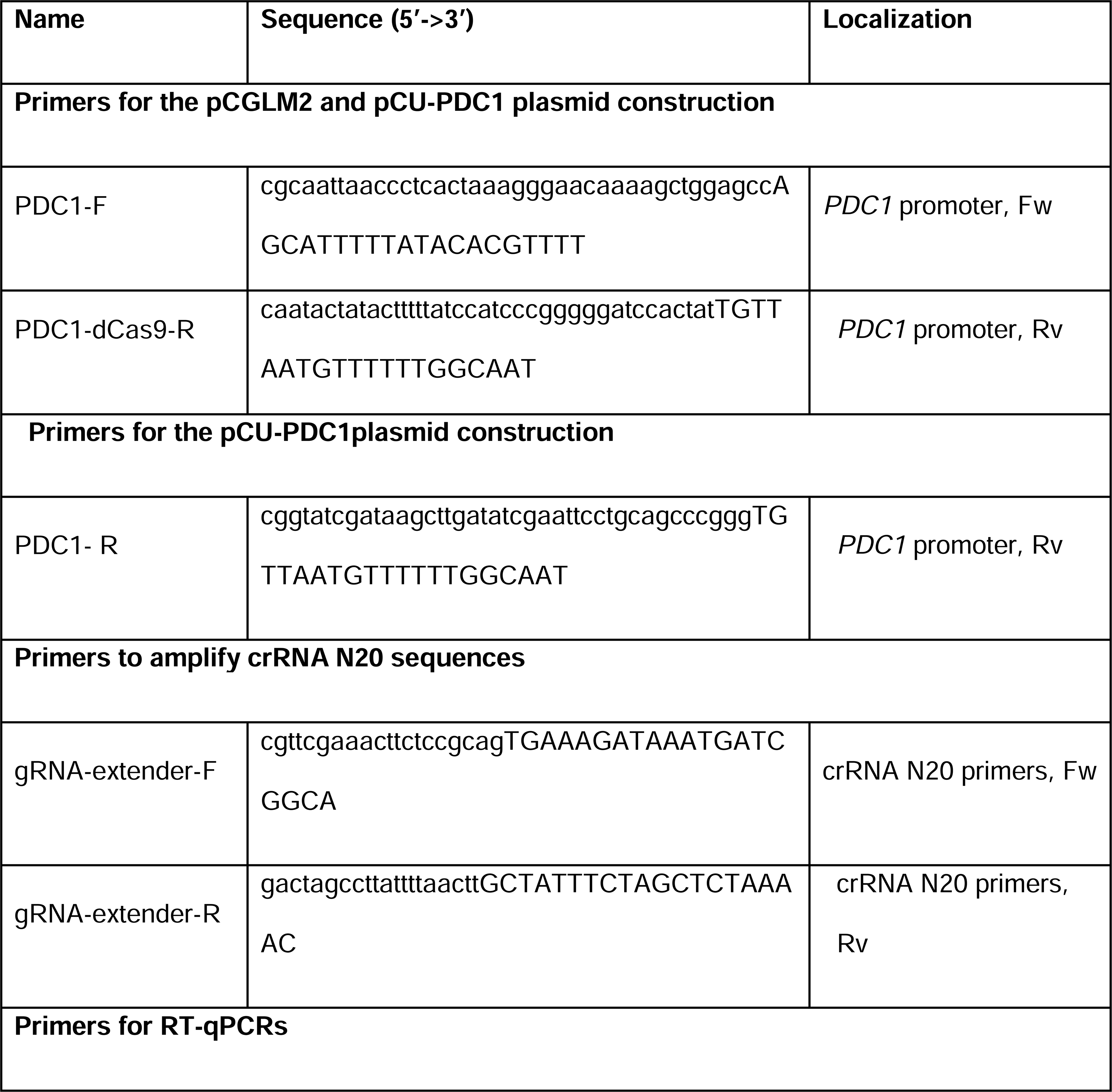
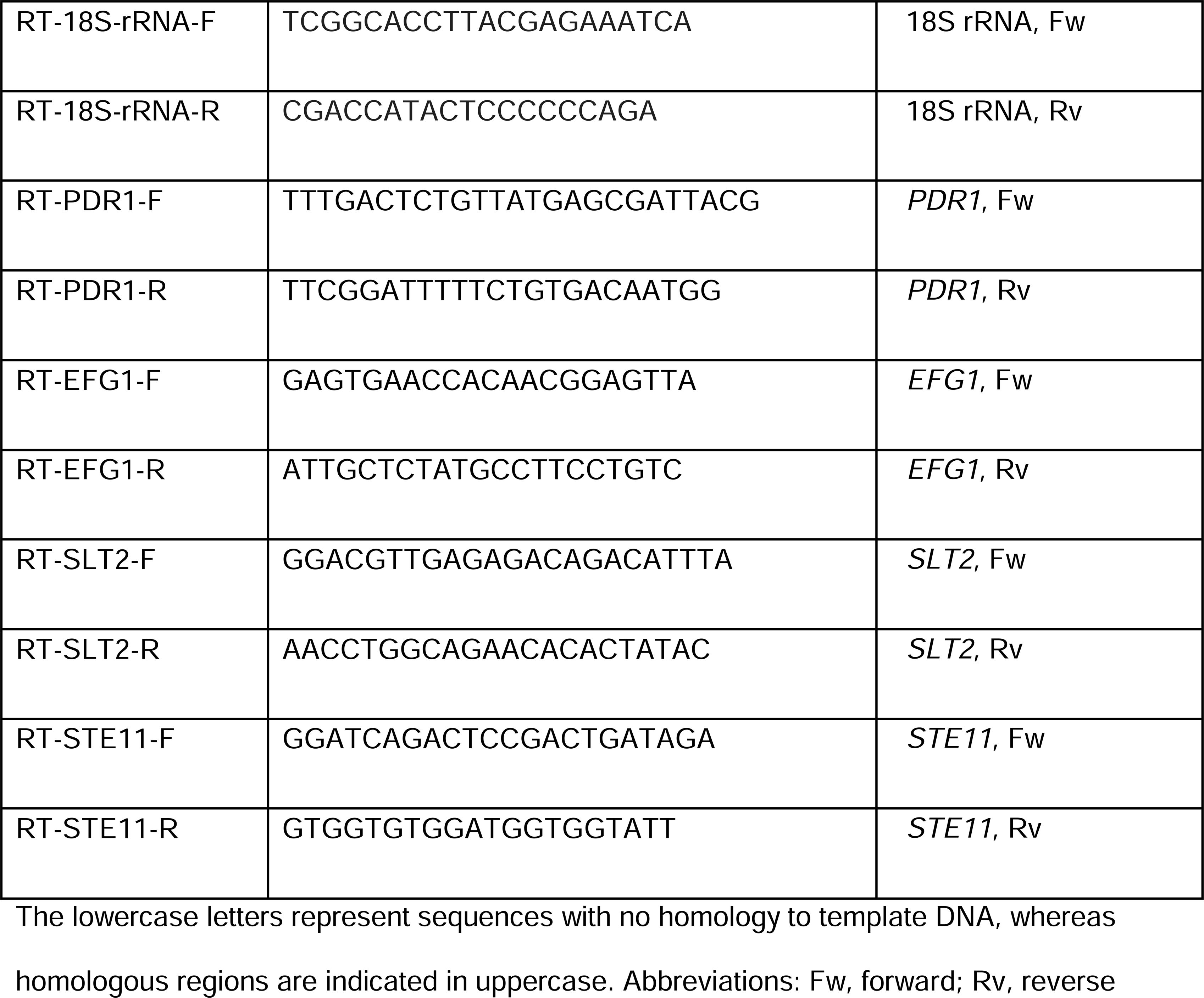
Primers used in this study.

**Supplementary Table 4.**
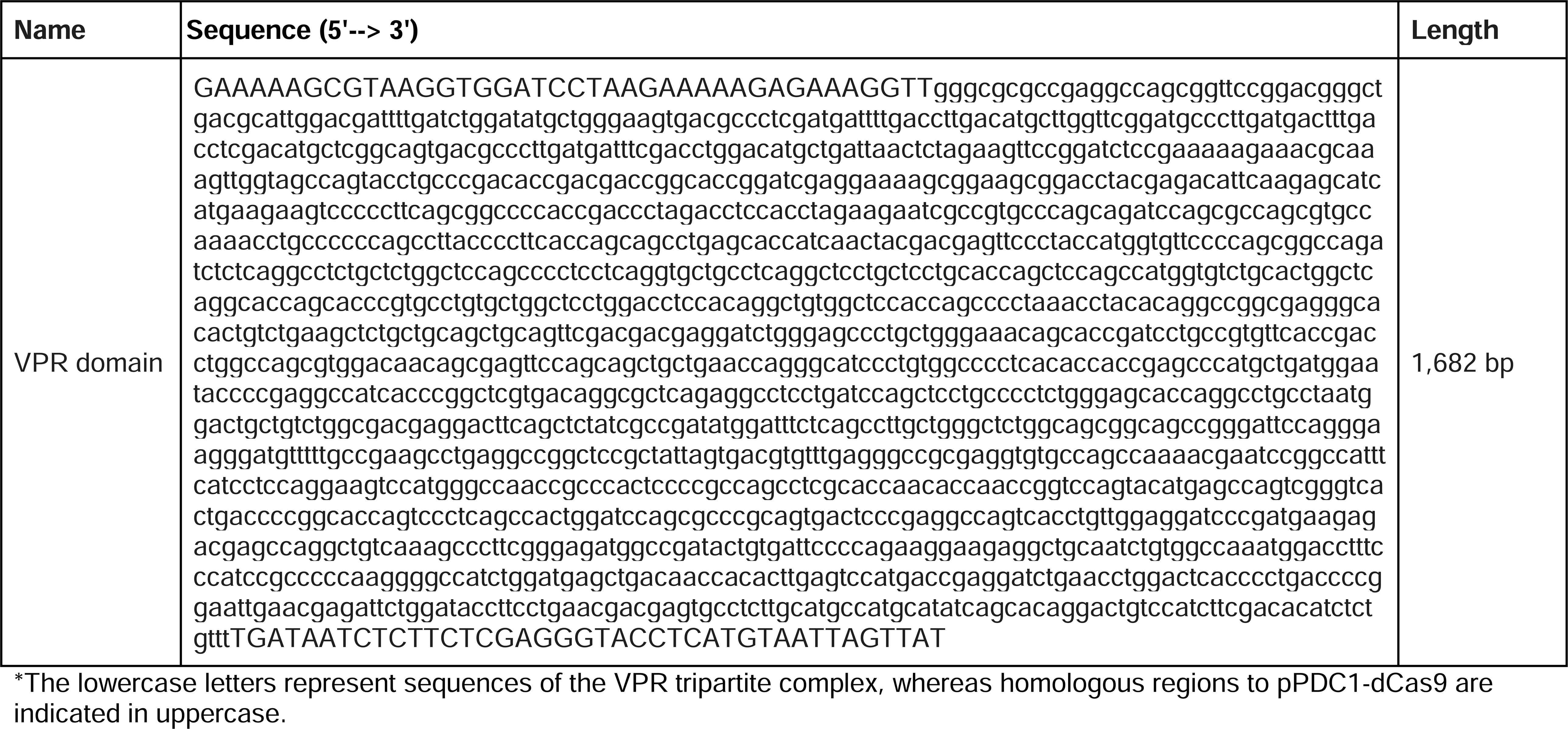
Gene fragments used in this study.

**Supplementary Table 5.**
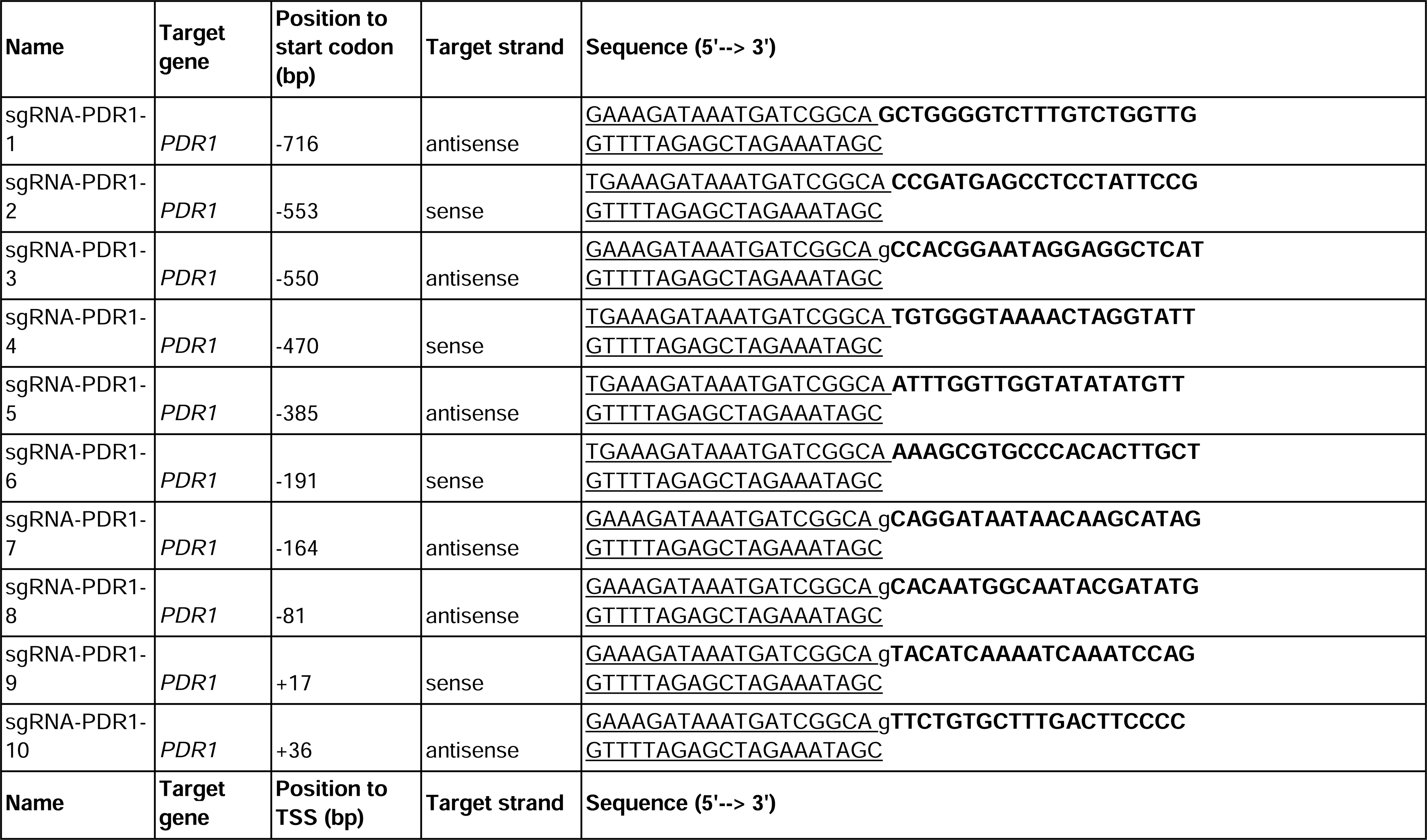
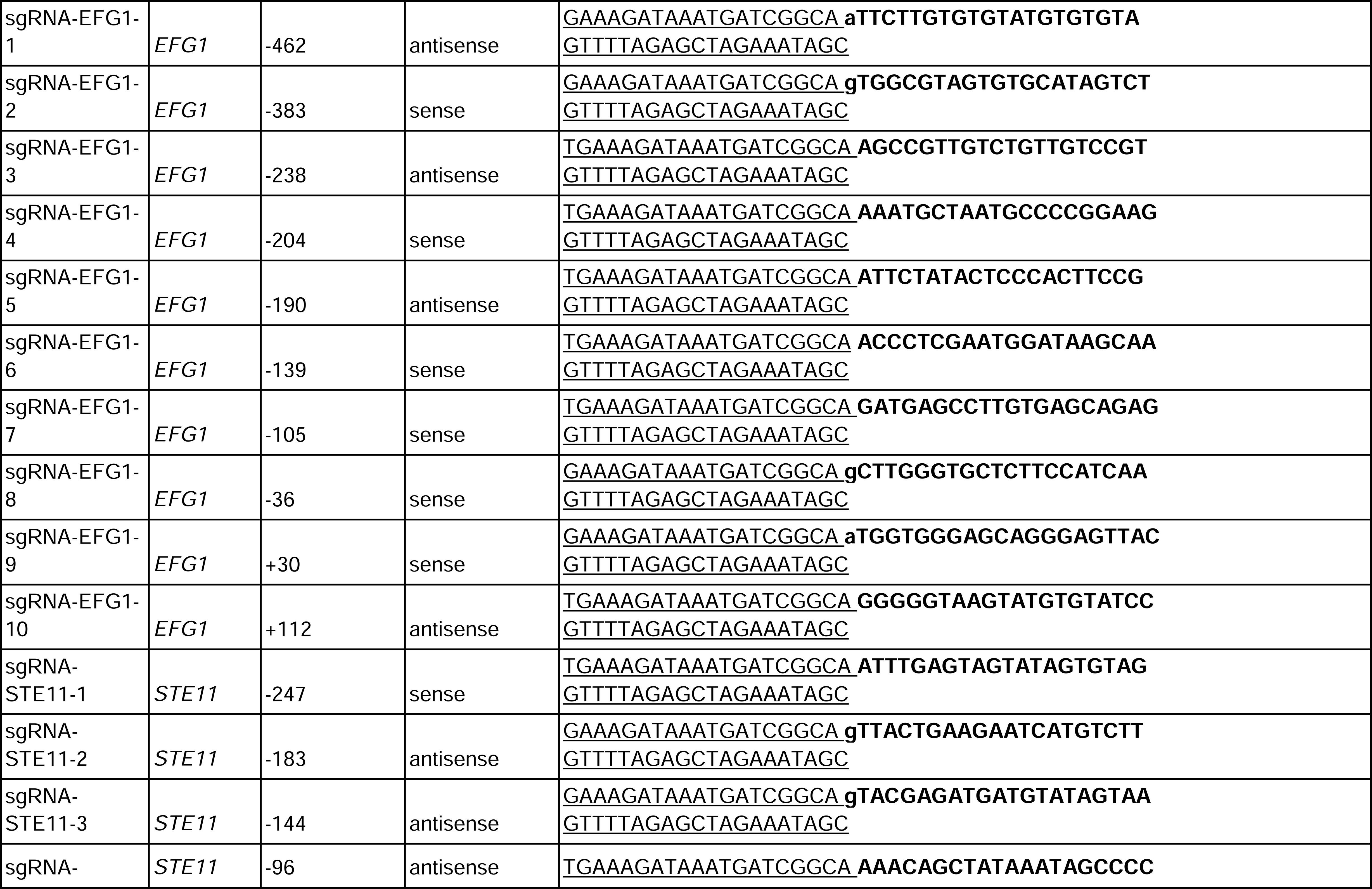
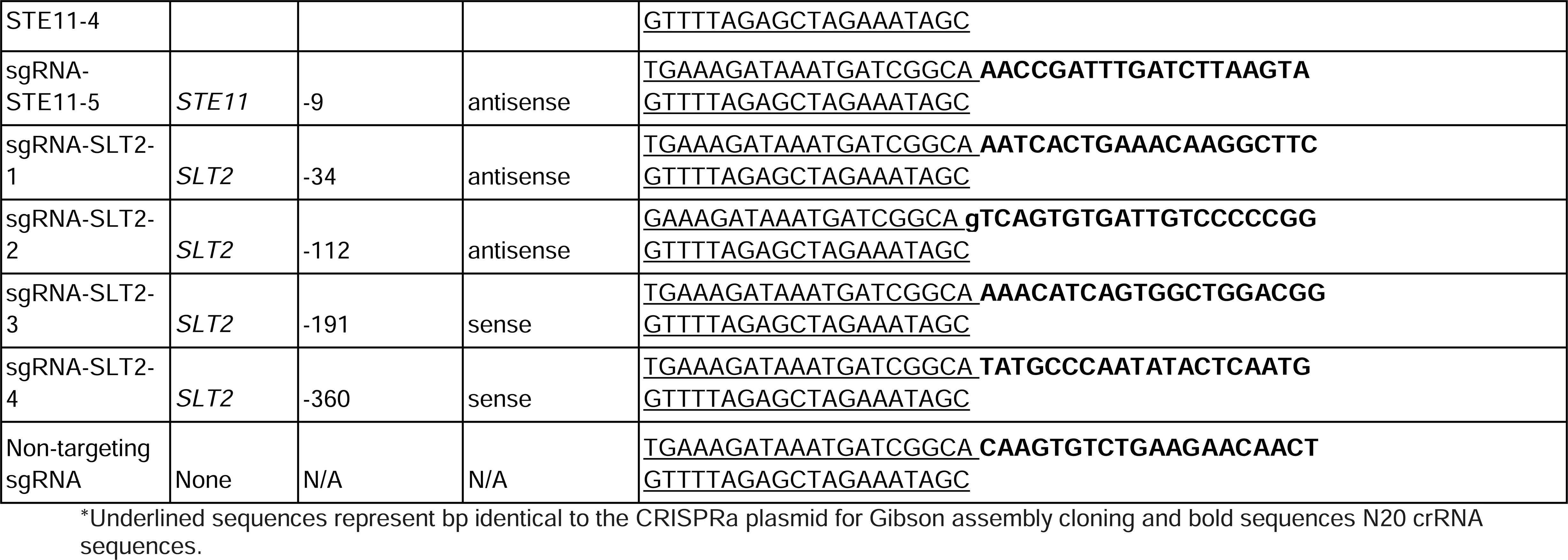
sgRNAs sequences used in this study.

## Notes

### Competing Interest Statement

The authors have declared no competing interest.

